# Highly efficient generation of isogenic pluripotent stem cell models using prime editing

**DOI:** 10.1101/2022.02.15.480601

**Authors:** Hanqin Li, Oriol Busquets, Yogendra Verma, Khaja Mohieddin Syed, Nitzan Kutnowski, Gabriella R. Pangilinan, Luke Gilbert, Helen Bateup, Donald C. Rio, Dirk Hockemeyer, Frank Soldner

## Abstract

The recent development of prime editing (PE) genome engineering technologies has the potential to significantly simplify the generation of human pluripotent stem cell (hPSC)-based disease models. PE is a multi-component editing system that uses a Cas9-nickase fused to a reverse transcriptase (nCas9-RT) and an extended PE guide RNA (pegRNA). Once reverse transcribed, the pegRNA extension functions as a repair template to introduce precise designer mutations at the target site. Here, we systematically compared the editing efficiencies of PE to conventional gene editing methods in hPSCs. This analysis revealed that PE is overall more efficient and precise than homology-directed repair (HDR) of site-specific nuclease-induced double-strand breaks (DSBs). Specifically, PE is more effective in generating heterozygous editing events to create autosomal dominant disease-associated mutations. By stably integrating the nCas9-RT into hPSCs we achieved editing efficiencies equal to those reported for cancer cells, suggesting that the expression of the PE components, rather than cell-intrinsic features, limit PE in hPSCs. To improve the efficiency of PE in hPSCs, we optimized the delivery modalities for the PE components. Delivery of the nCas9-RT as mRNA combined with synthetically generated chemically-modified pegRNAs and nicking guide RNAs (ngRNAs) improved editing efficiencies up to 13-fold compared to transfecting the prime editing components as plasmids or ribonucleoprotein particles (RNPs). Finally, we demonstrated that this mRNA-based delivery approach can be used repeatedly to yield editing efficiencies exceeding 60% and to correct or introduce familial mutations causing Parkinson’s disease in hPSCs.

## INTRODUCTION

One of the current challenges of using hPSCs to model human diseases is to precisely and efficiently engineer the genome to introduce designer mutations (Hockemeyer and Jaenisch, 2016; Soldner and Jaenisch, 2018). Currently, the predominant approach in hPSCs is to induce targeted double-strand DNA breaks (DSBs) using highly active site-specific nucleases, such as the CRISPR/Cas9 system (Cong et al., 2013; Ding et al., 2013; Jinek et al., 2012; Jinek et al., 2013; Mali et al., 2013) or protein engineering platforms including zinc finger nucleases (ZFNs) (Hockemeyer et al., 2009; Soldner et al., 2011; Zou et al., 2009) and transcription activator-like effector nucleases (TALEN) (Boch et al., 2009; Cermak et al., 2011; Hockemeyer et al., 2011). Such targeted DSBs have been shown to substantially increase genome editing efficiency over conventional homologous recombination. However, since nuclease-induced DSBs are in most context preferentially repaired by non-homologous end joining (NHEJ) compared to homology-directed repair (HDR) mechanisms, DSB-mediated genome editing frequently generates undesirable compound heterozygous editing outcomes with one correctly targeted allele and an insertion or deletion (indel) on the second allele, causing the disruption of the protein coding sequence (Cox et al., 2015). Therefore, it has been challenging to generate disease-associated dominant mutations in a heterozygous setting. By contrast, PE has been shown to overcome this limitation, as it does not require a DSB but directly repairs a nicked DNA strand (Anzalone et al., 2019). PE is a multi-component editing system composed of a Cas9-nickase fused to a reverse transcriptase (nCas9-RT) and an extended prime editing guide RNA (pegRNA) that is reverse transcribed and functions as a repair template at the target site (Anzalone et al., 2019). Here, we systematically compare different genome editing methods and show that PE is overall more efficient and precise to introduce heterozygous point mutations into hPSCs. Furthermore, by optimizing the delivery modality of the PE components, we were able to establish a highly efficient genome editing platform for hPSCs. By comparing plasmid, RNA-protein RNPs and *in vitro* transcribed mRNA delivery, we found that nucleofecting nCas9-RT as mRNA combined with synthetically generated and chemically-modified pegRNAs yielded editing efficiencies exceeding 60%, which is comparable to efficiencies observed in tumor cell lines (Anzalone et al., 2019; Nelson et al., 2021). Together, these data indicate that PE has the potential to greatly facilitate the generation of disease-specific hPSC models.

## RESULTS

To evaluate the potential use of PE to genetically modify hPSCs, we directly compared editing outcomes of PE to established CRISPR/Cas9 and TALEN targeting approaches with the goal of introducing disease-relevant point mutations. Initially, we chose to target the Leucine Rich Repeat Kinase 2 (*LRRK2*) gene to introduce the G2019S (G6055A) mutation (Gilks et al., 2005), which is one of the most frequent pathogenic substitutions linked to Parkinson’s disease (PD). This mutation is found in approximately 4% of dominantly inherited familial PD cases, in both heterozygous and homozygous forms and around 1% of sporadic PD cases (Healy et al., 2008). To introduce the G2019S (G6055A) mutation into hPSCs, we generated plasmid-based CRISPR/Cas9, TALEN and PE reagents (without [PE2] or with secondary ngRNA [PE3]) using previously established optimized design and targeting procedures (Figure 1A) (Anzalone et al., 2019; Hockemeyer et al., 2011; Hsu et al., 2021; Soldner et al., 2011; Soldner et al., 2016). Briefly, we co-electroporated the human embryonic stem cell (hESC) line WIBR3 with the respective genome engineering components and an enhanced green fluorescent protein (EGFP)-expressing plasmid to allow for the enrichment of transfected cells by fluorescence activated cell sorting (FACS). Genome editing outcomes were evaluated in the transfected bulk cell population (EGFP-positive) 72 hours after electroporation and in single cell-derived subclones. Next generation sequencing (NGS) of amplicons spanning the G2019S (G6055A) region indicated that 2% to 4% of alleles carried the correct G2019S (G6055A) substitution (Figures 1B and S1A). While we observed roughly comparable editing efficiencies using CRISPR/Cas9, TALEN and PE (with secondary ngRNA [PE3]) gene targeting, all HDR-based approaches generated a significantly higher number of undesired editing outcomes with 19.6% and 3.3% indels for CRISPR/Cas9 and TALEN, respectively, compared to less than 0.5% for PE (Figure 1B and S1A). Genotyping of expanded single cell-derived clones showed a higher efficiency in generating heterozygous correctly targeted cell lines using PE primarily due to a substantial higher number of compound heterozygous editing outcomes for CRISPR/Cas9 and TALEN targeting with the correctly inserted G6055A sequence variant on one allele and indels on the second allele (Figure 1C, as identified by RFLP and Sanger sequencing analysis). Together, these data indicate that PE is overall more efficient and substantially more precise in generating heterozygous mutations in hPSCs, as compared to traditional DSB-based genome engineering approaches.

**Figure 1.**
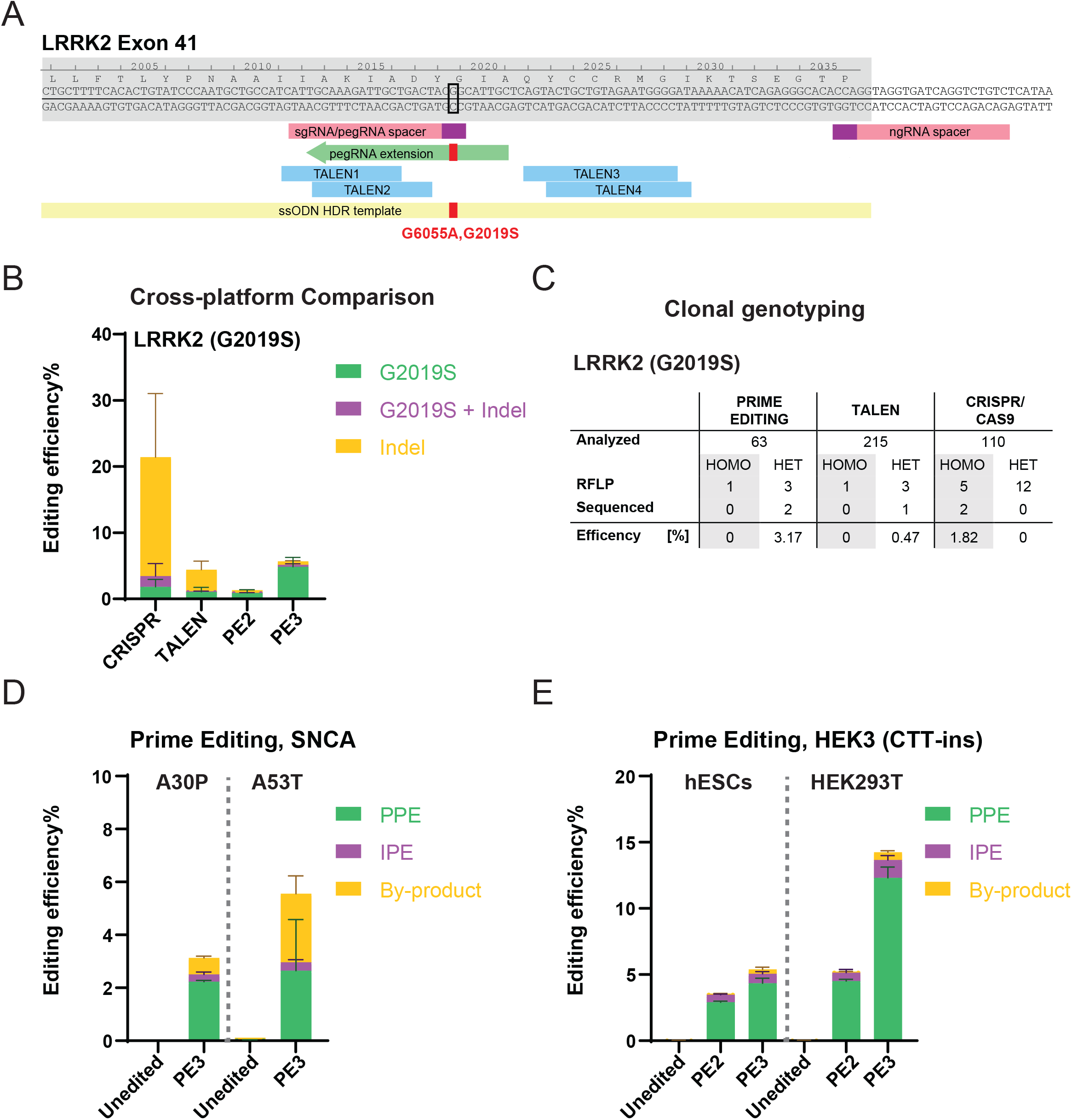
Systematic comparison of CRISPR/Cas9, TALEN and PE-based genome editing efficiencies in hESCs using plasmid-based delivery. **A.** Schematic depicting genome editing strategies to generate the *LRRK2* (G6055A, G2019S) mutation. Exon, grey shade; prospacers for CRISPR/Cas9 and PE, pink boxes; PAM sequences, purple boxes; representative pegRNA extension, green arrow; TALEN recognition sites, blue boxes; ssODN HDR template for CRISPR/Cas9 targeting, yellow box; the nucleotide to mutate, black open square; intended mutation, red filled square. **B.** Comparison of bulk genome editing outcomes between CRISPR/Cas9, TALEN and PE to insert the *LRRK2* (G2019S) mutation (aggregated analysis using four different pegRNA designs and two TALEN pairs), N=2 for CRISPR/Cas9, N=3 for TALEN, N=2 for PE2, N=6 for PE3. For individual analysis see Figure S1A. **C.** A summary of clonal genotyping comparing different genome editing strategies indicating the number and editing efficiency of single cell-derived clones carrying the correct heterozygous (HET) or homozygous (HOMO) G2019S substitution as identified by restriction fragment length polymorphism analysis (RLFP) followed by Sanger sequencing to excluded additional indels. **D.** Generating *SNCA* familiar PD mutations, A30P and A53T, using prime editing. PPE, precise prime editing; IPE, impure prime editing; By-product, other indels. A30P, N=3; A53T, N=2. **E.** Comparison of bulk prime editing outcomes on *HEK3* (CTT-insertion) edits between hESCs and HEK293T cells. Color scheme, same as Figure 1D. N=3. (Error bars indicate the standard deviation [SD], N = number of biological replicates).

To scale the PE-based genome editing approach and streamline the derivation of correctly modified single cell clones, we applied a recently established genome editing platform, which employs multiplex low cell number nucleofection, limited dilution, and NGS-dependent genotyping instead of the time consuming and laborious FACS sorting and manual single cell expansion steps to isolate correctly edited hPSC lines (see Figure S2 and Experimental Procedures for details). While this approach results in slightly lower overall bulk editing efficiencies (as determined by NGS), most likely due to the lack of FACS-based enrichment of transfected cells, the substantially reduced number of cells required and the streamlined workflow allows for highly efficient, multiplexed generation of genome-edited hPSC lines in parallel in less than 4 weeks (Figure S2). Importantly for this work, the omission of FACS enrichment allowed us to systematically and simultaneously compare a larger number of delivery modalities, as described below.

To confirm the feasibility of using PE to introduce point mutations efficiently and robustly into hPSCs, we tested PE at additional genomic loci. We were able to introduce mutations into the previously published and commonly targeted *HEK3* (CTT insertion) locus (Anzalone et al., 2019), as well as two additional PD-associated mutations into the *SNCA* (*α*-Synuclein) locus (A30P [G88C] (Krüger et al., 1998) and A53T [G209A] (Polymeropoulos et al., 1997); Figure S1B,C) with editing efficiencies comparable to the *LRRK2* locus (Figure 1D,E; quantified as pure prime editing efficiencies [PPE] as defined in Petri et al. (2021)). As described for *LRRK2*, the analysis of single cell-expanded clones revealed the efficient generation of heterozygous and homozygous hPSC lines carrying the dominant A30P mutation in the *SNCA* gene (Figure S3A,B). Importantly, representative cytogenetic analysis of single cell-expanded *LRRK2* (G2019S) and *SNCA* (A30P) clones showed normal karyotypes for 7 out of 7 tested cell lines. Together, these experiments demonstrate that PE can be used to robustly and efficiently introduce disease-associated mutations into hPSCs to generate isogenic disease models.

During these experiments, we noted that editing outcomes for both the PE2 and PE3 approaches appeared considerably lower than what was previously reported for a variety of human tumor cell lines (Anzalone et al., 2019; Nelson et al., 2021). Indeed, we found that plasmid-based targeting of the *HEK3* (CTT insertion) locus resulted in only ∼4.3% PPE in WIBR3 hESCs compared to ∼12.7% PPE in HEK293T tumor cells using the PE3 strategy (Figure 1E). Similar differences in gene editing efficiencies between hPSCs, primary cells and tumor cell lines have been commonly observed for other genome engineering approaches including CRISPR/Cas9 targeting (Bowden et al., 2020; Haapaniemi et al., 2018; Ihry et al., 2018). It remains unclear if this difference is the result of cell-intrinsic factors that restrict genome editing specifically in hPSCs or whether the low efficiency is a consequence of insufficient delivery of the PE components.

To test whether PE efficiencies could be increased by optimized delivery of the prime editor, we stably expressed the nCas9-RT protein (PE2 version of the prime editor protein as described in Anzalone et al. [2019]) followed by a 2A-EGFP fluorescent reporter from the *AAVS1* safe harbor locus (DeKelver et al., 2010; Hockemeyer et al., 2009) (Figure 2A). We established an hPSC clone that uniformly expressed GFP and maintained pluripotency (Figure 2B and S3C). Nucleofection of these cells with a plasmid encoding the *HEK3* (CTT insertion) pegRNA (without [PE2] or with [PE3] secondary ngRNA) or with a chemically-modified synthetic pegRNA (without [PE2] or with [PE3] secondary ngRNA) resulted in substantially increased editing efficiencies of up to 22% of correctly inserted modifications (Figure 2C). Similarly, targeting the *LRRK2* (G2019S) and *SNCA* (A30P) loci with chemically-modified synthetic pegRNAs (without [PE2] or with [PE3] secondary ngRNA) resulted in editing efficiencies up to 12% and 27%, respectively (Figure 2D). While these data do not exclude fundamental biological differences in the PE process between hPSCs and other cell types, these experiments demonstrate that the method of delivery of the PE components has a significant role in dictating the genome editing efficiency in hPSCs, which can be comparable to the efficiencies observed in tumor cell lines and primary cells.

**Figure 2.**
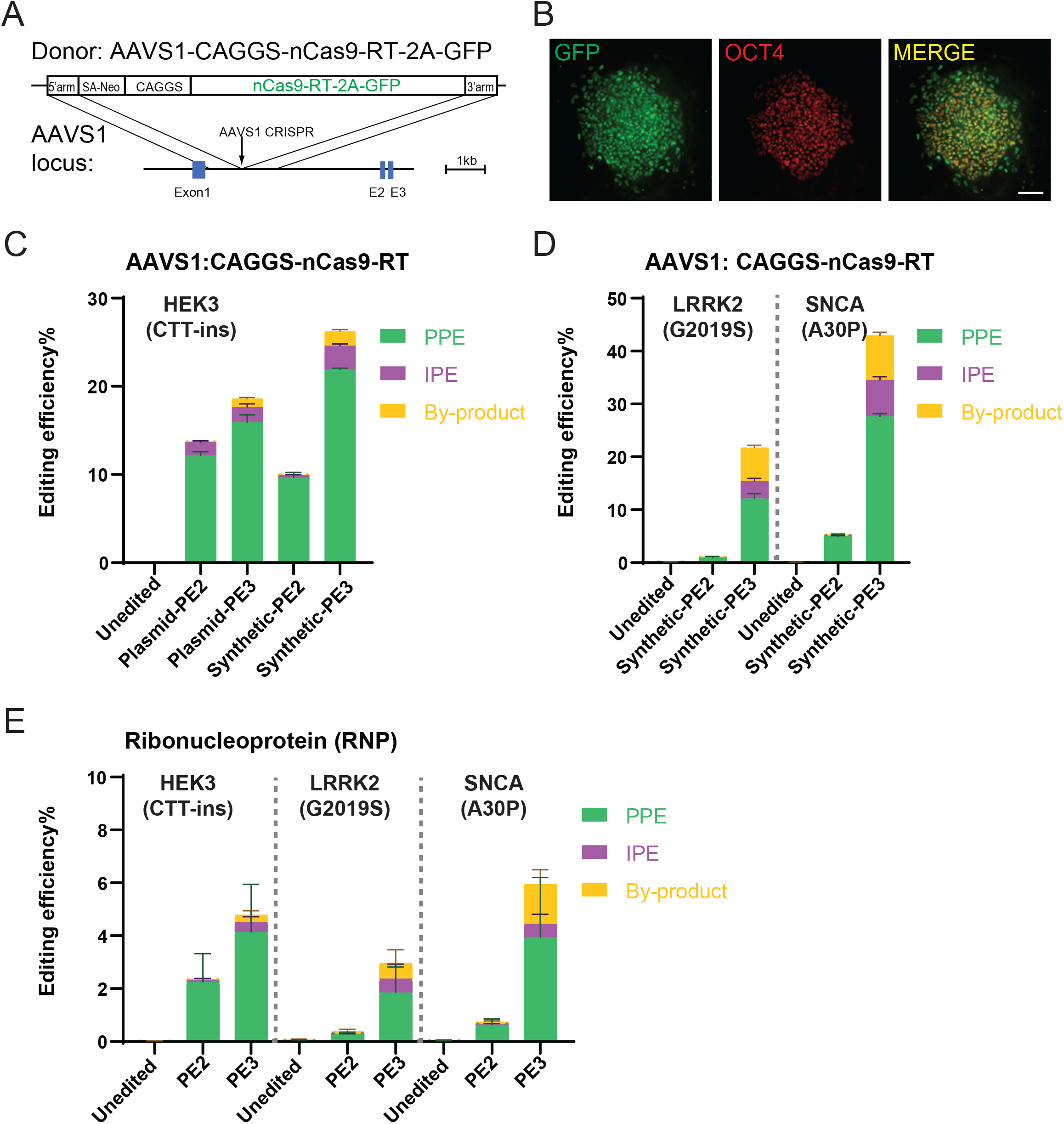
Prime editing in hESCs expressing nCas9-RT protein from the *AAVS1* safe harbor or delivered as RNP. **A.** Schematic of the genome editing strategy to knock-in nCas9-RT-2A-GFP (PE2 version of the prime editor protein as described in Anzalone et al. [2019]) into the *AAVS1* locus. **B.** Expression of GFP and immunostaining of OCT4 on hESCs with nCas9-RT-2A-GFP knock-in. Scale bar, 100µm **C.** Comparison of bulk prime editing outcomes between plasmid-expressed pegRNAs/ngRNAs and synthetic pegRNAs/ngRNAs on *HEK3* (CTT-insertion) edits in hESCs with nCas9-RT-2A-GFP knock-in. N=3. **D.** Bulk prime editing outcomes on *LRRK2* (G2019S) and *SNCA* (A30P) edits using synthetic pegRNAs/ngRNAs in hESCs with nCas9-RT-2A-GFP knock-in. N=3. **E.** Bulk prime editing outcomes from RNP delivery on *HEK3* (CTT-insertion), *LRRK2* (G2019S) and *SNCA* (A30P) edits in hPSCs. N=6. (Error bars indicate the standard deviation [SD], N = number of biological replicates).

To improve PE efficiencies using transient delivery of the PE components, we set out to optimize PE delivery conditions. Initially, we focused on delivering the PE components as RNPs, a highly efficient approach described for CRISPR/Cas9-mediated genome editing (Zuris et al., 2015), which was recently successfully adapted for PE in zebrafish and human primary T cells (Petri et al., 2021). Using recombinant nCas9-RT protein (PE2 version of the prime editor protein as described in Anzalone et al. [2019] purified from bacteria) (Figure S4A) and the previously established protocols for RNP-based CRISPR/Cas9 editing (Petri et al., 2021; Zuris et al., 2015), we nucleofected pre-assembled RNPs containing the recombinant nCas9-RT protein and chemically-modified synthetic pegRNAs (without [PE2] or with [PE3] secondary ngRNA) targeting the *HEK3* (CTT insertion), *LRRK2* (G2019S) and *SNCA* (A30P) loci. Consistent with previous reports in other cell types, we observed RNP-mediated editing outcomes in hPSCs with locus-dependent efficiencies between 1% and 6% (Figure 2E and S4B,C). While these data clearly indicate the feasibility of RNP-based PE in hPSCs, the observed efficiencies are comparable to the plasmid-based approach and far below the efficiencies observed with stable expression of nCas9-RT from the *AAVS1* locus. To exclude that RNP-based PE efficiencies were concentration- or batch-dependent, we repeated some of these experiments with protein from independently purified nCas9-RT batches (Figure S4D) and used higher protein concentrations (Figure S4E). However, none of these conditions resulted in substantially improved RNP-based editing efficiencies in hPSCs (Figure S4D,E).

An alternative approach, allowing highly efficient delivery of Cas9 for CRISPR/Cas9-based genome editing (Chang et al., 2013; Hwang et al., 2013; Wang et al., 2013), is to deliver the prime editor using *in vitro* transcribed mRNA (Chen et al., 2021; Surun et al., 2020). To systematically compare plasmid-, RNP- and mRNA-based PE at the above established *HEK3* (CTT insertion), *LRRK2* (G2019S) and *SNCA* (A30P) loci, we nucleofected either: *(i)* plasmid-based nCas9-RT together with plasmid-based pegRNAs (without [PE2] or with [PE3] secondary ngRNA), *(ii)* preassembled RNPs containing nCas9-RT protein and chemically-modified synthetic pegRNAs (without [PE2] or with [PE3] secondary ngRNA) or *(iii) in vitro* transcribed nCas9-RT mRNA together with chemically-modified synthetic pegRNAs (without [PE2] or with [PE3] secondary ngRNA). Bulk NGS revealed that the combination of *in vitro* transcribed mRNA-based delivery of the nCas9-RT with chemically-modified synthetic pegRNAs and ngRNAs consistently increased editing efficiencies across all three tested loci up to 13-fold compared to plasmid- and 8-fold compared to RNP-delivery (Figure 3A). Using this optimized *in vitro* transcribed mRNA-based delivery approach allowed us to achieve editing efficiencies up to 26.7% (for the *SNCA* locus). Surprisingly, when combined with secondary nicking of the non-edited strand (PE3), these editing efficiencies were even higher than using the stably nCas9-RT expressing cell lines and comparable to efficiencies commonly observed in human tumor cell lines (Anzalone et al., 2019; Nelson et al., 2021). While mRNA-based editing efficiencies seem to depend on the approach used to *in vitro* transcribe the nCas9-RT mRNA (Figure S5A), we found that mRNA-based editing efficiencies are highly consistent across different nCas9-RT mRNA batches when using the best *in vitro* transcription conditions (Figure S5B). Furthermore, we observed that using single-stranded mRNA compared to either RNPs or double-stranded plasmid DNAs resulted in improved overall health and increased survival of single cells as indicated by increased clonal survival following nucleofection (Figure 3B). To test if the high efficiency of mRNA-based PE is a unique feature of the WIBR3 hESCs, we repeated some key experiments in a second human-induced pluripotent stem cell (hiPSC) line 8858, (Pasca et al., 2015) by targeting the *LRRK2* (G2019S) and *SNCA* (A30P) loci and found comparable editing efficiencies (Figure 3C). Importantly, we were able to establish single cell-derived clones carrying the correct *SNCA* (A30P) and *LRRK2* (G2019S) mutations with high efficiency (Figure 3D). Considering that potential therapeutic applications would require precise genome-editing of hPSCs in xeno-free conditions, we were able to show efficient and robust PE of the *LRRK2* (G2019S) locus in WIBR3 hESCs using several commonly used feeder-free culture systems with comparably high PE efficiencies (Figure S5C).

**Figure 3.**
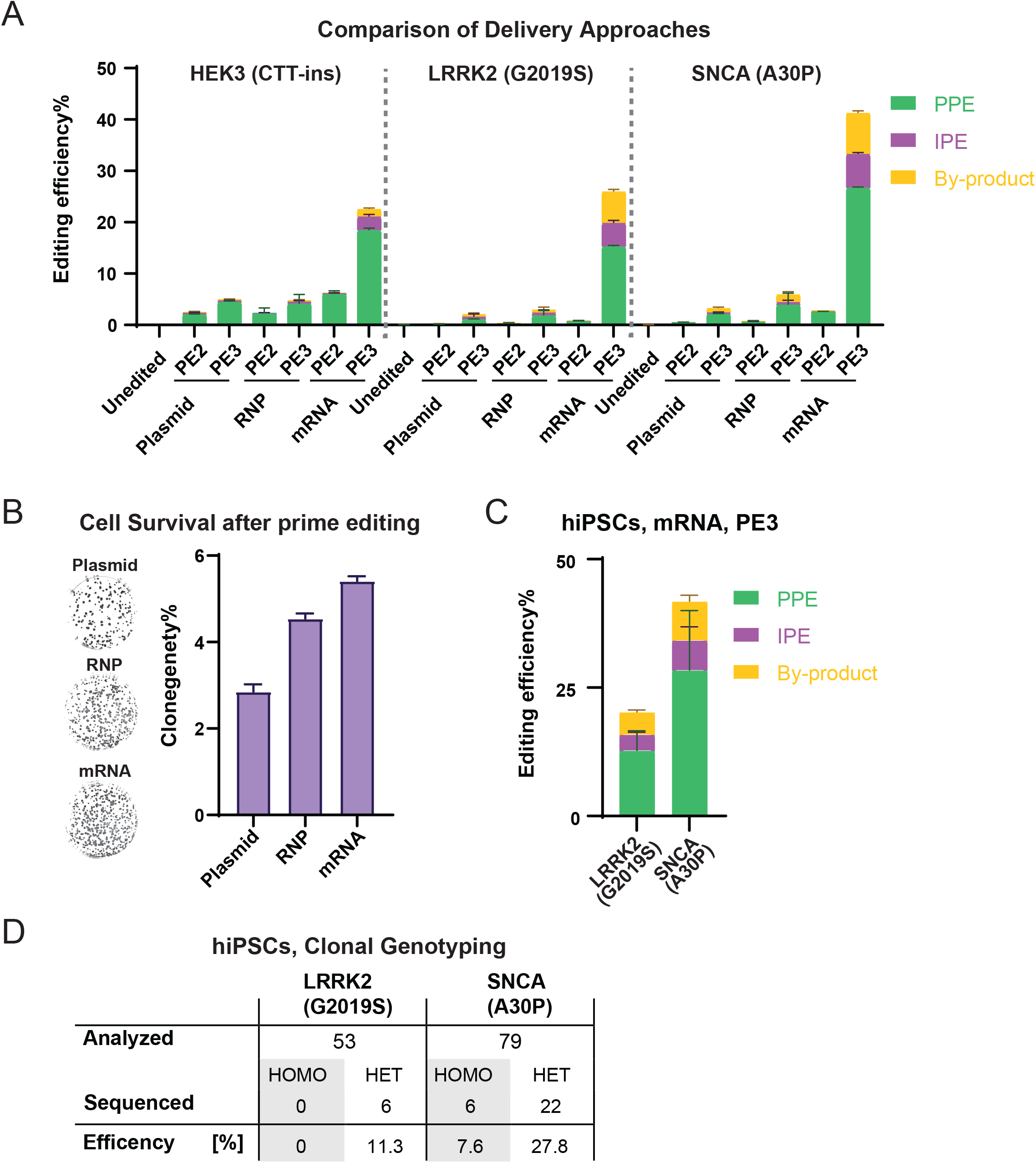
Highly efficient prime editing in hPSCs using mRNA-based delivery. **A.** Comparison of bulk prime editing outcomes between plasmid, RNP and mRNA-based delivery on indicated modifications in hESCs. Plasmid, mRNA groups, N=3; RNP data shown in Figure 2E was included in this analysis for direct comparison, N=6. **B.** Representative images and quantification of alkaline phosphatase staining comparing clonogenicity of hESCs after nucleofection between plasmid, RNP and mRNA-based delivery. N=2. **C.** Bulk prime editing outcomes on *LRRK2* (G2019S) and *SNCA* (A30P) edits in a hiPSCs line using mRNA-based delivery. N=2. **D.** A summary of clonal genotyping from *LRRK2* (G2019S) and *SNCA* (A30P) prime editing in hiPSCs indicating the number and editing efficiency of single cell-derived clones carrying the correct heterozygous (HET) or homozygous (HOMO) substitution. (Error bars indicate the standard deviation [SD], N = number of biological replicates).

A major limitation of classical CRISPR/Cas9 targeting remains the high number and complexity of undesirable editing outcomes (indels). These alleles are resistant to targeting with the same reagents and thus limit the overall HR-editing efficiencies in the context of continued editing or retargeting. Given the much-reduced occurrence of indel-containing alleles in mRNA-based PE, we hypothesized that this approach might allow efficient retargeting of the same locus. Indeed, we find for all tested loci (*HEK3*, *LRRK2*, *SNCA*) that additional rounds of mRNA-based prime editing of the same cell population could substatially increase overall editing efficiencies (Figure 4A). This indicates that multiple rounds of mRNA-based PE could result in precise and nearly complete editing of a bulk hPSC population without any type of selection.

**Figure 4.**
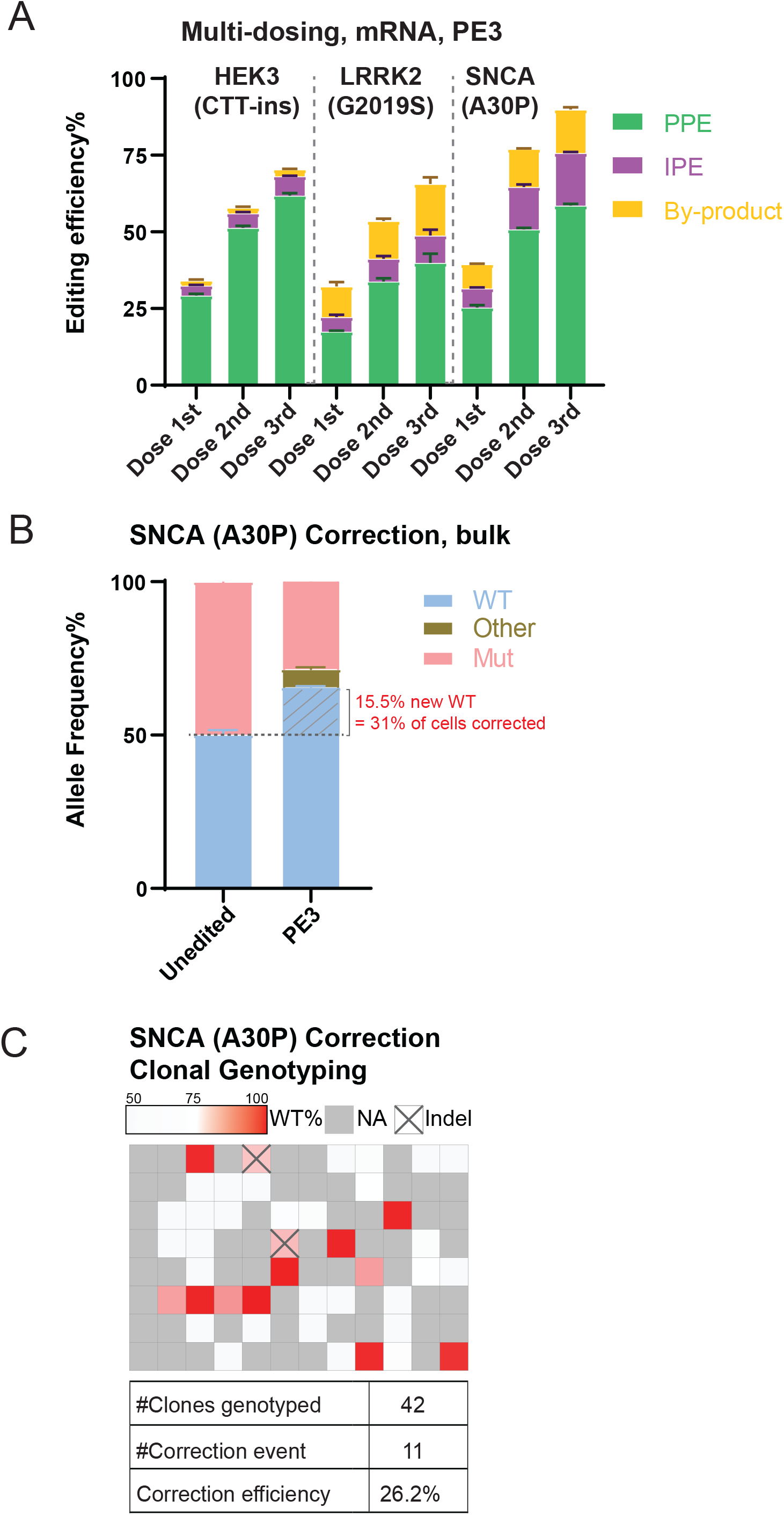
Repeated prime editing and reversion of an *SNCA* (A30P) mutation in hESCs. **A.** Comparison of bulk prime editing outcomes in a multi-dosing strategy using mRNA-based delivery on the three indicated mutations in hESCs. N=3. **B.** Bulk NGS analysis indicating allele spectrum before (unedited) and after mRNA-bases prime editing (PE3) to correct the heterozygous *SNCA* (A30P) mutation in hESCs. N=3. The dash line indicates the 50% allele frequency. The portion of WT allele converted from the mutant allele is hatched. **C.** Heatmap and summary of clonal genotyping from a *SNCA* (A30P) heterozygous mutation correction experiment in a 96-well format. The wells containing more than 5% indel reads were labeled as Indel. NA indicates wells that without hESCs. (Error bars indicate the standard deviation [SD], N = number of biological replicates).

The data presented thus far describes the insertion of disease-associated mutations into a wild-type genetic background. To test whether mRNA-based PE can be used to correct disease-causing mutations, we designed pegRNAs to specifically target only the mutated *SNCA* (A30P) allele to revert this mutation back to wild-type in hPSCs. When targeting a heterozygous *SNCA* (A30P) hESC line, bulk NGS indicated the correction of 31.0% of the mutated A30P alleles (Figure 4B). Subsequent genotyping of single-cell derived clones indicated 26.2% precisely corrected clones without additional undesired modifications (indels) of the wild-type allele (Figure 4C). Taken together, our data indicate that *in vitro* transcribed mRNA-based PE is a highly efficient gene editing approach in hPSCs that has the potential to greatly facilitate the generation of disease-specific hPSC models.

## DISCUSSION

The experiments performed here provide a detailed experimental road map for how to implement PE towards genome engineering of hPSCs. We show that mRNA transfection of the prime editor component (nCas9-RT) paired with the transfection of chemically-modified guide RNAs is well tolerated and highly effective for introducing precise designer mutations in hiPSCs and hESCs. Considering that mRNA-based PE does not require specialized molecular or biochemical skills and consistently achieves high editing efficiency in hESC and hiPSC lines, we predict that this approach has the potential to greatly facilitate the generation of disease-specific hPSC models and will be widely adopted by researchers.

During the process of establishing this workflow, we made several key observations. We find that PE can be as efficient in hPSCs as has been reported for cancer cells (Anzalone et al., 2019; Nelson et al., 2021). We demonstrate that this approach efficiently allows to introduce or correct heterozygous disease-related mutations in hPSCs with base pair precision and without introducing undesired additional modifications on the second allele. The resulting cells showed a normal karyotype, consistent with low genotoxicity of PE due to the lack of DSBs (Anzalone et al., 2019). In the past, generating such isogenic sets cells that differ exclusively at individual disease-causing sequence variants was highly laborious and an experimental bottleneck. Here we overcome this challenge by deploying PE via optimized delivery methods. We demonstrate that hPSCs can be subjected to several rounds of PE, eventually yielding up to 60% correctly targeted alleles. Importantly, PE efficiencies might be further increased by including mRNAs coding for DNA mismatch repair inhibiting proteins, a novel approach that has been recently shown to significantly improve the prime editing platform (Chen et al., 2021). These very high editing efficiencies without the need for selection of enrichment of targeted clones provide an intriguing platform to develop more robust *in vitro* disease models and potential therapeutic applications of PE in hPSCs or differentiated cell types.

## Limitations of the study

In our study we successfully introduced three out of three familial PD point mutations into hPSCs using previously established algorithms to design peRNAs (Hsu et al., 2021). In each case a classical protospacer adjacent motif (PAM) was present close to the intended amino acid substitution. Our work did not explore more complex or challenging genetic modifications, however we expect that systematic approaches that establish optimized design parameters for prime editing, as recently described for cancer cells (Kim et al., 2020; Nelson et al., 2021) and the development of Cas9 variants with non-classical PAMs (Chatterjee et al., 2020; Kleinstiver et al., 2015; Miller et al., 2020) combined with the optimized protocols reported here will allow PE to become a general method of choice for genome editing in hPSCs.

## EXPERIMENTAL PROCEDURES

### hPSCs culture

All hESC and hiPSC lines were routinely maintained on irradiated or mitomycin C-inactivated mouse embryonic fibroblast (MEF) feeder layers as described previously (Soldner et al., 2016). Detailed protocols for culturing of MEFs and hPSCs can be found on protocols.io (dx.doi.org/10.17504/protocols.io.b4msqu6e; dx.doi.org/10.17504/protocols.io.b4pbqvin). The hiPSC 8858 line (Sergiu Pasca lab, Stanford) (Pasca et al., 2015) and hESC line WIBR3 (NIH Registration Number: 0079; Whitehead Institute Center for Human Stem Cell Research, Cambridge, MA) (Lengner et al., 2010) were maintained on MEFs in hESC media (DMEM/F12 [Thermo Fisher Scientific] supplemented with 15% fetal bovine serum [FBS] [Hyclone], 5% KnockOut Serum Replacement [Thermo Fisher Scientific], 1 mM glutamine [Invitrogen], 1% nonessential amino acids [Thermo Fisher Scientific], 0.1 mM β-mercaptoethanol [Sigma] and 4 ng/mL FGF2 [Thermo Fisher Scientific /Peprotech])], 1X Penicillin & Streptomycin [Thermo Fisher Scientific]. Cultures were passaged every 5 to 7 days with collagenase type IV (Invitrogen; 1 mg/mL). The identities of all parental hESC and hiPSC lines were confirmed by DNA fingerprinting and all cell lines were regularly tested to exclude mycoplasma contaminations using a PCR-based assay. If required (as indicated for the methods for the respective experiments) hiPSCs were adapted to feeder-free conditions on Vitronectin (VTN-N, Thermo Fisher Scientific) coated plates in mTeSR-plus (StemCell Technologies) or StemFlex (Thermo Fisher Scientific) media. Detailed protocols feeder-free culturing of hPSCs can be found on protocols.io (dx.doi.org/10.17504/protocols.io.b4mcqu2w).

### Culturing and transfection of HEK293T cells

HEK393T cells were maintained in HEK293T media (DMEM [Thermo Fisher Scientific], 15% FB Essence [Avantor], 2mM glutamine [Thermo Fisher Scientific], 1mM nonessential amino acids [Thermo Fisher Scientific], 1X Penicillin & Streptomycin [Thermo Fisher Scientific]), and passaged every other day with 0.25% Trypsin with EDTA [Thermo Fisher Scientific]). For transfection, cells were seeded into 0.2% gelatin-coated 12-well plates at 1×10^4^ cell/cm^2^. One day later, cells in each well were transfected with 500ng pCMV-PE2 (Anzalone et al., 2009, Addgene#132775), 330ng pU6-pegRNA and 170ng pBPK1520-ngRNA for PE3 strategy or 500ng pCMV-PE2, 500ng pU6-pegRNA for PE2 strategy using 1Dl lipofectamine 2000 [Thermo Fisher Scientific] in opti-MEM [Thermo Fisher Scientific]. Cells were collected for genomic DNA extraction and NGS-based allele quantification 3 days post transfection.

### Molecular cloning

PegRNAs-expressing plasmids (pU6-pegRNA) were cloned by ligating annealed oligo pairs (Supplemental table 2) with BsaI-digested pU6-peg-GG-acceptor (Addgene #132777) as described previously (Anzalone et al., 2019). CRISPR-RNA expressing plasmids (px330-GFP) targeting the *LRRK2* locus were cloned by ligating annealed oligo pairs (Supplemental table 2) with BbsI-digested px330-GFP as described previously (Soldner et al., 2016). Nicking ngRNA-expressing plasmids (pBPK1520-ngRNA) were cloned by ligating annealed oligo pairs (Supplemental table 2) with BsmBI-digested pBPK1520 (addgene#65777) as described previously (Kleinstiver et al 2015). The heterodimeric TALEN pairs to target *LRRK2* (G2019S) were constructed and tested as described previously (Cermak et al., 2011; Hockemeyer et al., 2011) using the following RVD sequences. Pair#1: TALEN1 (Plasmid LRRK2-TALEN-TA01L): 5’-HD-NI-NG-NG-NN-HD-NI-NI-NI-NN-NI-NG-NG-NN-HD-NG-3’ and TALEN3 (Plasmid LRRK2-TALEN-TA03R): 5’-HD-HD-HD-HD-NI-NG-NG-HD-NG-NI-HD-NI-NN-HD-NI-NN-NG-NI-HD-NG-3’. Pair#2: TALEN2 (Plasmid LRRK2-TALEN-TA04L): 5’-NN-HD-NI-NI-NI-NN-NI-NG-NG-NN-HD-NG-NN-NI-NG-3’ and TALEN4 (Plasmid LRRK2-TALEN-TA07R): 5’-NI-NG-HD-HD-HD-HD-NI-NG-NG-HD-NG-NI-HD-NI-NN-HD-NI-NN-NG-3’.

### mRNA *in vitro* transcription

The plasmid pCMV-PE2 was cleaved with restriction endonuclease PmeI (100µg DNA in 1mL) for 4 hr. at 37°C. The cleaved DNA was isolated by phenol-chloroform extraction and ethanol precipitation and resuspended at 500µg/mL in TE buffer. The DNA was stored at −20°C. Eight 20µL *in vitro* transcription reactions were set up using 1µg of template DNA in each reaction using the New England Biolabs HiScribe T7 ARCA kit with tailing (E2060S) (as per the manufacturer’s instructions) and incubated for 2 hr. at 37°C in an incubator (not a temp block). Using eight 20µL reactions, after transcription, DNase I treatment and polyA tailing (as per the manufacturer’s instructions), the RNA was purified on four 50μg New England Biolabs Monarch RNA cleanup columns (T2040L) (as per the manufacturer’s instructions) and eluted in 25μL per column RNase-free H_2_O and pooled. The RNA was stored at −80°C and yield from the total of eight reactions was ∼200μg purified PE2 mRNA by measuring A_260_ on a Nanodrop 2000c spectrophotometer. Note that the nCas9-RT fusion mRNA is ∼ 6,500 nt. Detailed protocols can be found on protocols.io (dx.doi.org/10.17504/protocols.io.b3fmqjk6).

### nCas9-RT protein purification

The nCas9-RT fragment from pCMV-PE2 was retrieved by BglII digestion then cloned into the pET30a(+) expression vector in frame between the NotI and NdeI sites using NEBuilder® HiFi DNA Assembly Master Mix (NEB) with bridging gblocks (Supplemental Table 1) to encode a version of the protein bearing a C-terminal His6-tag. For protein expression, the plasmid was introduced into Rosetta 2 (pLysS). The cells were grown at 37°C and shaken at 175 rpm. IPTG (0.5 mM) was added at an OD600 of 0.6 and the cells were grown for 16 hr. at 18°C. The cells were harvested by centrifugation (5000 x g) for 10 min. at 4°C. Harvested cell pellets were washed with PBS and snap-frozen in liquid nitrogen for later purification. Cell pellets were thawed on ice, disrupted in 35mL lysis buffer (25mM HEPES-KOH pH 7.6, 1M KCl, 20mM imidazole, 1mM DTT, 1mM PMSF and 1X protease inhibitor cocktail), briefly sonicated, then clarified by centrifugation at 25,000xg for 30 min. Supernatants were filtered through a 0.22μm syringe filter before application to 5mL of nickel resin (Ni-NTA Superflow, QIAGEN) equilibrated in loading buffer (25mM HEPES-KOH pH 7.6, 150mM KCl, 20mM imidazole, 1mM DTT and 1mM PMSF). The resin was washed with 100mL loading buffer followed by 50mL wash buffer (25mM HEPES-KOH, pH 7.6, 150mM KCl, 40mM imidazole, 1mM DTT and 1mM PMSF). Protein was eluted in batch six times with 10mL elution buffer (wash buffer + 500mM imidazole). The eluted protein was diluted into a low-salt buffer (25mM HEPES-KOH pH 7.6, 100mM KCl, 1mM DTT and 1mM PMSF), then loaded onto a 1mL HiTrap heparin HP column (GE Healthcare) pre-equilibrated in low salt buffer and eluted with a linear gradient of 100mM to 1M KCl over 40 column volumes (CVs). Peak fractions were concentrated to 8mg/mL using a Spin-X UF 20 50kDa MWCO (Corning). Protein concentration was determined by UV absorbance at wavelength of 280nm. The final protein purity was determined by SDS–PAGE and Coomassie staining to be around 90%. The C-terminal His6-tag was not removed prior to the experiments. Detailed protocols can be found on protocols.io dx.doi.org/10.17504/protocols.io.b4yxqxxn).

### Prime editing, CRISPR/Cas9 and TALEN-based genome editing using plasmid vectors

As indicated for the respective experiments, plasmid vector-based prime editing was performed using electroporation or nucleofection (using the high throughput hPSCs genome editing pipeline described below). Detailed protocols can be found on protocols.io (dx.doi.org/10.17504/protocols.io.b4qnqvve).

Briefly for electroporation-based plasmid-mediated TALEN, CRISPR/Cas9 and prime editing, hPSCs were cultured on MEFs in Rho-associated protein kinase (ROCK)-inhibitor (10μM, Stemgent; Y-27632) for 24 hr. before electroporation. Cells were harvested using 0.05% trypsin/EDTA solution (Thermo Fisher Scientific) and resuspended in phosphate buffered saline (PBS). 1 × 10^7^ cells were electroporated (Gene Pulser Xcell System, Bio-Rad: 250 V, 500 mF, 0.4cm cuvettes) with the following plasmid vectors: For the prime editing PE2 strategy, we used 33µg pCMV-PE2-GFP ((Anzalone et al., 2019), Addgene#132776) and 12µg pU6-pegRNA. For the prime editing PE2 strategy, we used 33µg pCMV-PE2-GFP, 12µg pU6-pegRNA and 5µg pBPK1520-ngRNA. For TALEN editing, we used 7.5µg for each (left and right) TALEN-nuclease plasmid, 10µg pEGFP-N1 and 26µg ssODN (single strand oligonucleotide containing the respective modification). For CRISPR/Cas9 editing, we used 16µg pX330-GFP gRNA and 26µg ssODN (single strand oligonucleotide containing the respective modification). A list of the respective plasmids can be found in Supplemental table 2. Cells were maintained on MEFs for 72 hr. in the presence of ROCK inhibitor followed by FACS sorting (FACS-Aria; BD-Biosciences) of a single-cell suspension. EGFP expressing cells were either were either directly used for bulk NGS based allele quantification or subsequently plated at a low density on MEFS in hESC media supplemented with ROCK inhibitor for the first 24 hr. Individual colonies were picked and expanded 10 to 14 days after electroporation. Correctly targeted clones were subsequently identified by RFLP and genomic sequencing (See Supplemental table 1 for respective primer sequences).

For nucleofection-based prime editing, hPSCs were cultured on MEFs in ROCK inhibitor for 24 hr. before nucleofection. Cells were harvested using collagenase IV (Thermo Fisher Scientific) followed by accutase (Thermo Fisher Scientific) and 5X10^5^ cell were resuspended in 20µL nucleofection solution and nucleofected (4D-Nucleofector TM Core + X Unit [Lonza], nucleofection program P3 primary cell, CA137) using the following plasmids vectors: For the prime editing PE2 strategy, we used 500ng pCMV-PE2 and 500ng pU6-pegRNA. For the prime editing PE2 strategy, we used 500ng pCMV-PE2, 330ng pU6-pegRNA and 170ng pBPK1520-ngRNA. A list of the respective plasmids can be found in Supplemental table 2. After nucleofection, cells were maintained either on MEFs in hESC media or on VTN-N coated plates in feeder-free media, both containing Rock Inhibitor and either used for NGS-based allele quantification or single cell cloning (following the high throughput hPSCs genome editing pipeline described below).

### Prime editing using RNP

Human PSCs cultured on MEFs were harvested and nucleofected using the same procedure as described in the plasmid delivery section, except with RNPs consisting of 90pmol purified nCas9-RT protein, 300pmol chemically modified synthetic pegRNA (Supplemental table 2) for PE2 strategy or 90pmol purified nCas9-RT protein, 200pmol chemically-modified synthetic pegRNA and 100pmol chemically modified synthetic ngRNA (Supplemental table 2) for PE3 strategy. For increased protein doses, 270pmol purified nCas9-RT protein were used instead. All RNPs were pre-assembled at room temperature for 10 min before nucleofection. Detailed protocols can be found on protocols.io. (dx.doi.org/10.17504/protocols.io.b4qnqvve).

### Prime editing using mRNA

Human PSCs cultured on MEFs were harvested and nucleofected using the same procedure as described in the plasmid delivery section, except with 4μg *in vitro* transcribed nCas9-RT mRNA, 150pmol chemically modified synthetic pegRNA for PE2 strategy or 4μg *in vitro* transcribed nCas9-RT mRNA, 100pmol chemically-modified synthetic pegRNA and 50pmol chemically modified synthetic ngRNA for PE3 strategy. For feeder feel culture, hPSCs were harvested using accutase. In multi-dosing experiments, after the 1st nucleofection, hPSCs were nucleofected for the 2nd and 3rd time at day 7 and day 14, respectively. Detailed protocols can be found on protocols.io (dx.doi.org/10.17504/protocols.io.b4qnqvve).

### Genotyping of single cell expanded genome edited hPSCs clones by RLFP

Restriction fragment length polymorphism (RFLP) analysis was performed as previously described (Hernandez et al., 2005) to screen single cell expanded clones for the insertion of the *LRRK2* (G2019S) mutation. Genomic DNA was amplified with primers SP-LRRK2-RLFP and ASP-LRRK2-RLFP (Supplemental table 1) under standard PCR conditions followed by restriction digest with SfcI. The *LRRK2* (G6055A, G2019S) mutation at coding nucleotide 6055 of *LRRK2* creates a novel SfcI cleavage site which allows to distinguish the two alleles after separation on a 3% agarose gel with the reference allele generating fragments of 228bp and 109bp and the mutated allele generating fragments of 207bp, 109bp and 21bp. Mutation carrying clones were further analyzed using Sanger and NGS sequencing of the PCR product.

### High throughput hPSCs genome editing pipeline

After nucleofection, cells were directly seeded onto MEF 96-well plates, at seeding densities of 10 cells/well in hPSCs media containing Rock inhibitor. Media was changed at day 4, 7, 10, 12, 13 and Rock inhibitor was supplemented at day 13. At day 14, cells were washed with PBS once then treated with 40µL 0.25% trypsin for 5 min. at 37°C, then 60µL hPSC media containing Rock inhibitor was added to each well to inactivate trypsin. Cells were then gently triturated and 50 µL cell suspension was transferred to a 96-well PCR plate pre-loaded with 50µL 2X lysis buffer (100mM KCl, 4mM MgCl_2_, 0.9% NP-40, 0.9% Tween-20, 500µg/mL proteinase K, in 20mM Tris-HCl, pH 8) for DNA extraction. The remaining 50µL of cells were reseeded to a new MEF 96-well plate preloaded with 100µL hPSC media containing Rock inhibitor and cultured for another 7 days with hPSC media changed daily. Meanwhile, the lysed cells in 96-well plates were incubated at 50°C overnight and then heated to 95°C for 10 min. to inactivate the proteinase K. A ∼300bp genomic region covering the designed mutation were amplified using primers (Supplemental table 1) containing NGS barcode attachment sites (GCTCTTCCGATCT) from 2ul cell lysis from each well with Titan DNA polymerase. Amplicons were purified at the UC Berkeley DNA Sequencing Facility, then i5/i7 barcoded in indexing PCR, pooled and sequenced on 150PE iSeq in the NGS core facility at the Innovative Genomics Institute (IGI). CRISPResso2 (Clement et al., 2019) in prime editing mode was used to analyze the NGS data to identify wells containing the designed mutation, with the following criteria. Heterozygous candidates: number of reads aligned >100, 70% >mutant allele frequency >20%, indels frequency <5%; homozygous candidates: number of reads aligned >100, mutant allele frequency >70%, indels frequency <5%. Cells in those identified wells were single cell subcloned once to ensure clonality. Detailed protocols for high throughput hPSCs genome editing (dx.doi.org/10.17504/protocols.io.b4mmqu46) and genotyping by next generation sequencing (dx.doi.org/10.17504/protocols.io.b4n3qvgn) can be found on protocols.io.

### *AAVS1* locus knock-in

To clone the AAVS1-SA-neo-CAGGS-nCas9-RT-2A-GFP targeting vector, the PmeI/SacII digested nCas9-RT fragment of pCMV-PE2-GFP were Gibson assembled into the EcoRI/KpnI sites of a parental AAVS1-SA-neo-CAGGS vector (a gift from Dr. John Boyle). Human PSCs cultured on MEFs were harvest and nucleofected as described in the plasmid delivery section, except with 1ug targeting vector and pre-assembled RNP consisting of 80pmol purified Cas9 (Macrolab, UC Berkely) and 300pmol chemically-modified sgRNA (Synthego) targeting the *AAVS1* locus. Cells were replated onto DR4 MEFs post nucleofection in hESC media containing Rock Inhibitor then selected with 70μg/mLG418 (invitrogen) for 10 days with media change daily from day 3. Survived clones were manually picked, expanded, gDNA extracted and PCR genotyped with primers (Supplemental table 1) flanking each homologous arms using PrimeStar GXL DNA polymerase (Takara). Correctly targeted clones were further expanded and banked. Detailed protocols can be found on protocols.io (dx.doi.org/10.17504/protocols.io.b37kqrkw).

### Karyotyping using aCGH

Human hPSCs cultured on MEFs were harvested using collagenase IV as big aggregates and settled 3 times in washing media (DMEM [Thermo Fisher Scientific], 5% Newborn Calf Serum [Sigma], 1X Penicillin & Streptomycin [Thermo Fisher Scientific]), then strained by an 80μm strainer. Cell aggregates that did not pass through the strainer were collected, snap frozen as cell pellet, then sent to Cell Line Genetics (Madison, WI) for aCGH karyotyping.

### Pluripotent marker staining

For Immunostaining, hPSCs cultured on MEFs were fixed in 4% paraformaldehyde at room temperature (RT) for 10 min., permeabilized in 0.3% Triton-X100/PBS for 20 min., blocked in blocking solution (3% BSA/PBS) for 1hr, incubated with primary antibody (OCT4 [PCRP-POU5F1-1D2, DSHB], 1:200; SSEA4 [MC-813-70, DSHB], 1:200) in blocking solution at 4°C overnight, then washed with PBS and incubated with secondary antibody [Thermo Fisher A11001, 1:1000] in blocking solution at room temperature for 1hr. For alkaline phosphatase staining, cells were fixed with cold 4% paraformaldehyde for 10 minutes, equilibrated with 100mM Tris-HCl, pH9.5 for 10 min. at RT, then incubated with NBT/BCIP (SK-5400, Vector laboratories) at room temperature for 2hr. to overnight. Images were acquired on a Zeiss Axio Observer A1 inverted fluorescence microscope. Detailed protocols can be found on protocols.io (dx.doi.org/10.17504/protocols.io.b4yyqxxw).

### Single cell survival assay

Human PSCs nucleofected with plasmid, RNP or mRNA were seeded to MEFs at 100 cells/cm^2^, then cultured for 14 days with media changed every other day. Cells were then stained for alkaline phosphatase as described above and the number of colonies in each condition were counted.

### Bulk NGS and allele quantification

Edited bulk cells were collected using trypsin at days 5 post nucleofection, then DNA extracted, mutation-region amplified, NGS and analyzed as described above Detailed protocols can be found on protocols.io (dx.doi.org/10.17504/protocols.io.b4n3qvgn). From the CRISPResso2 reported “Quantification_of_editing_frequency” table, the allele frequency of each group was calculated as following:

Wild type (WT), ([Unmodified Reference] + [Only Substitution Reference]) / [Total Reads aligned]

Pure primed editing (PPE), ([Unmodified Prime-edited] + [Only substitution Prime-edited]) / [Total Reads aligned]

Impure primed editing (IPE), ([Total Prime-edited]- [Unmodified Prime-edited] – [Only substitution Prime-edited]) / [Total Reads aligned]

By-product, 1-WT-PPE-IPE

### Software and Statistics

Bar graphs were drawn in Graphpad Prism 9. Error bars indicate the standard deviation (SD). Number of biological replicates (N) is indicated in each figure legend. Heatmaps were generated using Morpheus (Broad Institute).

## Acknowledgements

We would like to thank Devin Snyder for administrative support and help with editing of the manuscript. We thank Steven Poser for technical support, help with human hPSC culture and molecular biology. We thank all the members of the Hockemeyer, Soldner, Rio, Gilbert and Bateup lab for helpful discussions and comments on the manuscript. We thank Sergiu P. Pașca for providing the 8858 hiPSC line. This research was funded in part by Aligning Science Across Parkinson’s ASAP-000486 through the Michael J. Fox Foundation for Parkinson’s Research (MJFF). Some of the Flow Cytometry and Genomics shared resources at Albert Einstein College of Medicine were supported by the Cancer Center Support Grant (P30 CA013330).). H.L. is a fellow in the Siebel Stem Cell Institute. F.S. is supported by internal research support from the Department of Neuroscience at Albert Einstein College of Medicine. For the purpose of open access, the author has applied a CC BY public copyright license to all Author Accepted Manuscripts arising from this submission.

## Author contributions

H.L., O.B, L.G., H.S.B., D.R., D.H., and F.S designed the experiments. H.L. and O.B performed the editing experiments with the help of Y.V., K.H., and G.R. N.K. purified the recombinant PE-protein. H.L., O.B, D.R. D.H., and F.S. analyzed data and wrote the paper with the help of all other authors.

## Declaration of interests

The authors declare no competing interests.

## Data and Materials Availability

All sequencing data will be deposited in publicly accessible databases. Plasmids referred to in this paper have been deposited to Addgene’s Michael J. Fox Foundation Plasmid Resource and their associated doi can be found in Supplemental table 2.

## SUPPLEMANTAL FIGURE LEGENDS

**Figure S1.**
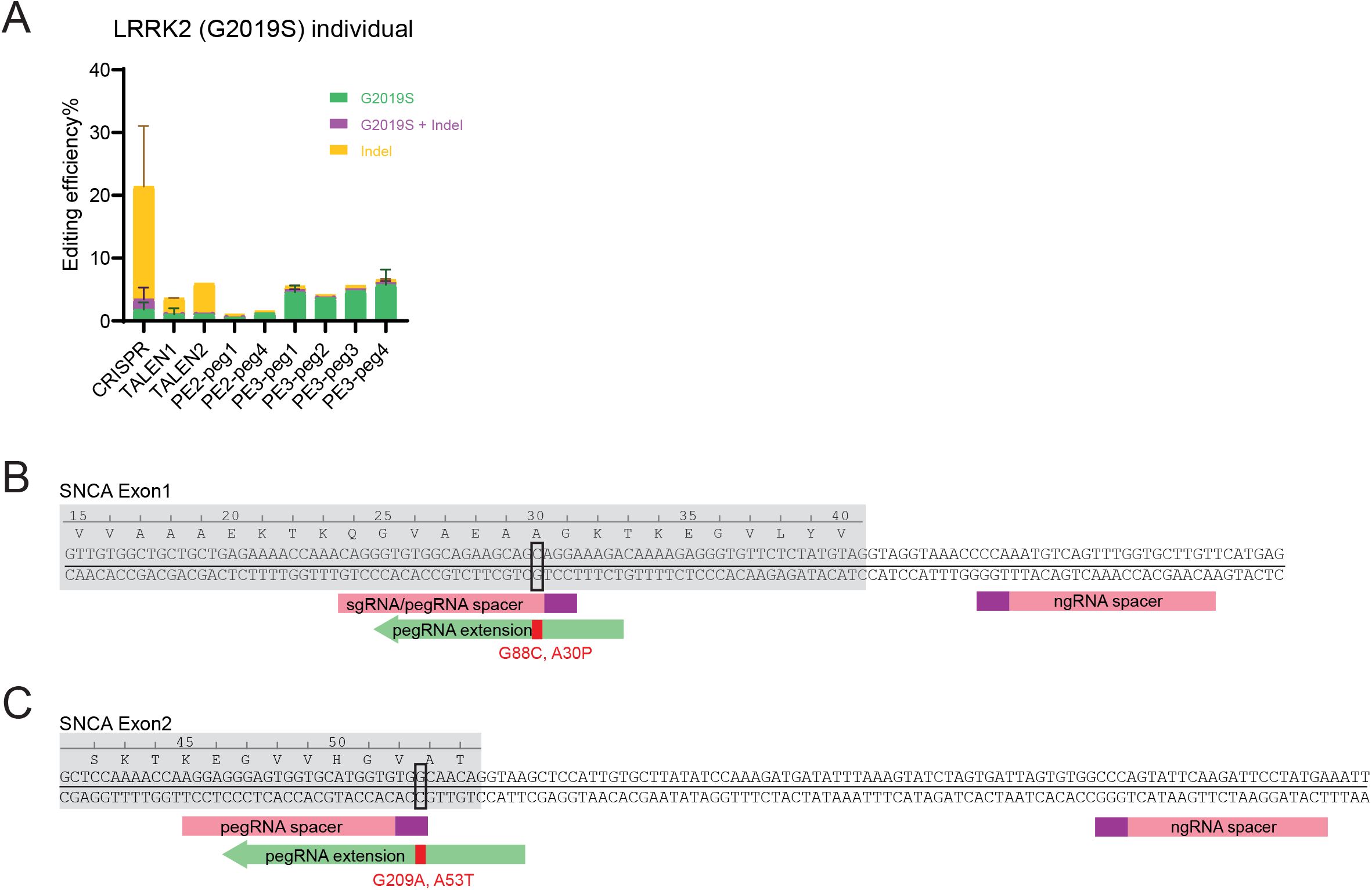
Schematics of *SNCA* PD mutation prime editing strategies and detailed *LRRK2* genome editing outcomes. **A.** Comparison of bulk genome editing outcomes between CRISPR, two TALEN pairs and four prime editing designs on *LRRK2* (G2019S). Individual analysis of aggregated data displayed in Figure 1B. CRISPR, N=2; TALEN pair#1, N=2; TALEN pair#2, N=1; PE2-peg1, N=1; PE2-peg4, N=1; PE3-peg1, N=2; PE3-peg2, N=1; PE3-peg3, N=1; PE3-peg4, N=2. (Error bars indicate the standard deviation [SD], N = number of biological replicates). **B.** Schematic of prime editing strategy for *SNCA* (A30P) mutation. Exon, grey shade; prospacers in prime editing, pink boxes; PAM sequences, purple boxes; pegRNA extension, green arrow. **C.** Schematic of prime editing strategy for *SNCA* (A53T) mutation. Color scheme, same as in B.

**Figure S2.**
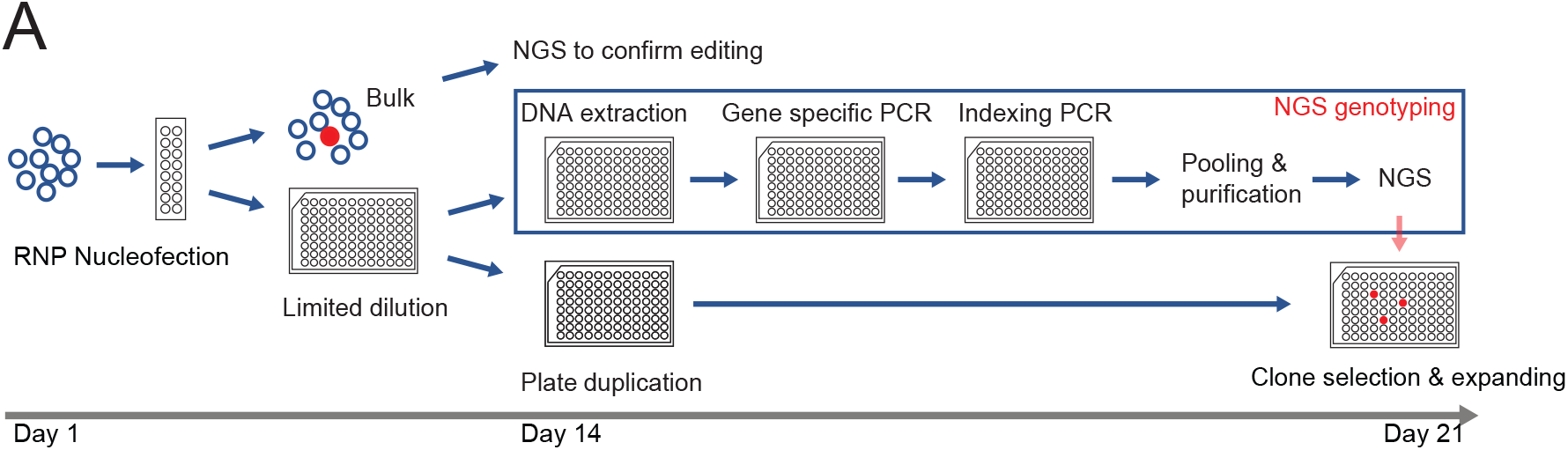
A high throughput hPSCs genome editing pipeline combining limited dilution and NGS genotyping. Nucleofected hPSCs are directly seeded into 96-well plates by limiting dilution. Upon colonies growing up, plates are duplicated to allocate one plate for cell maintainance and one plate for NGS genotyping. Clones with the desired genotype are expanded to establish genome edited hPSCs line.

**Figure S3.**
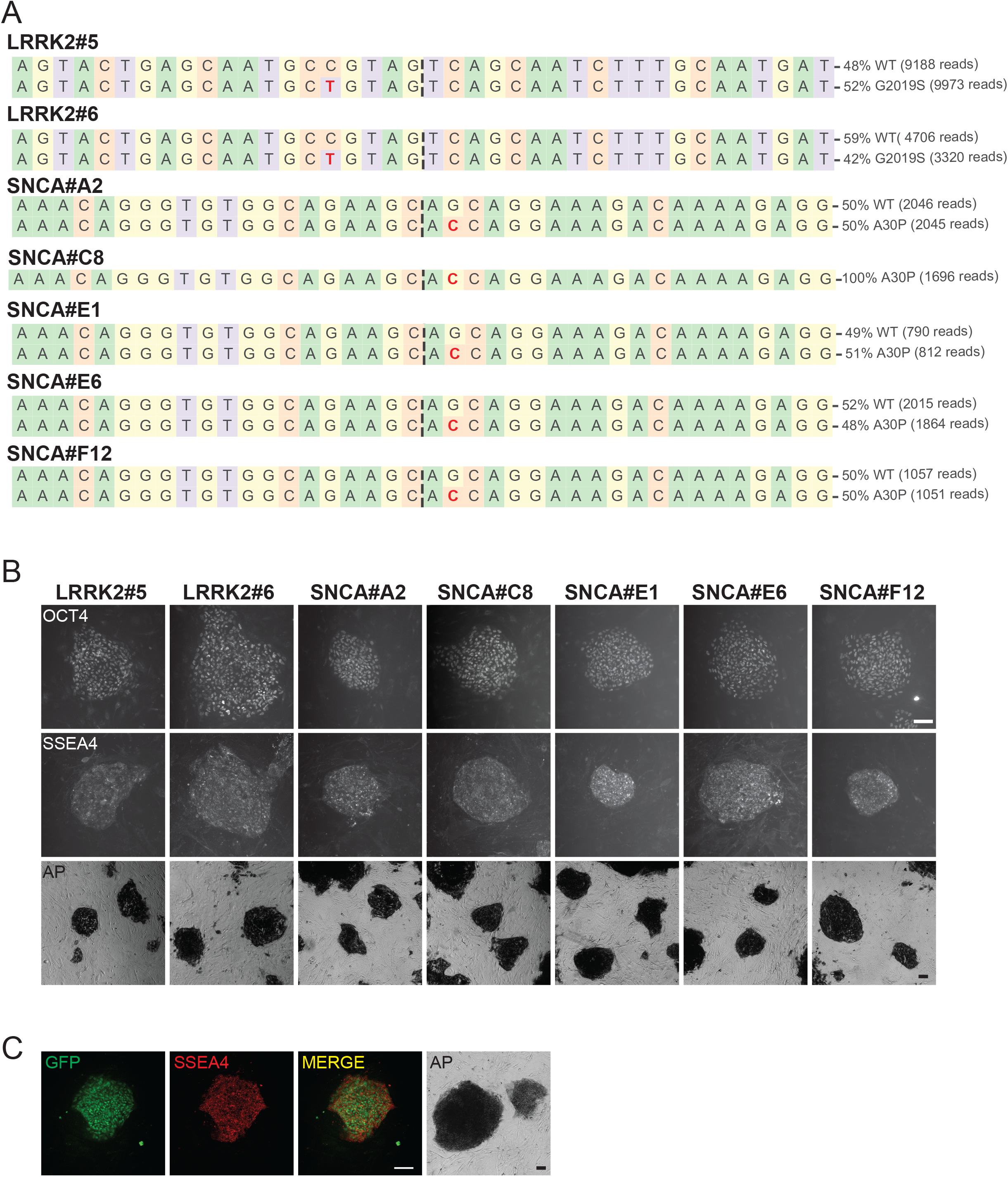
Genotyping and pluripotent marker staining of genome edited hESCs lines. **A.** NGS genotyping of the established hESCs lines harboring *LRRK2* (G2019S) and *SNCA* (A30P) mutations, introduced by prime editing with plasmid-based delivery. The mutated nucleotides are highlighted in red. **B.** Pluripotent marker staining of the edited hESCs lines shown in A. for OCT4, SSEA4 and alkaline phosphatase (AP). Scale bar, 100µm. **C.** GFP expression, SSEA4 immunostaining and alkaline phosphatase staining on hESCs with nCas9-RT-2A-GFP knock-in. Scale bar, 100µm.

**Figure S4.**
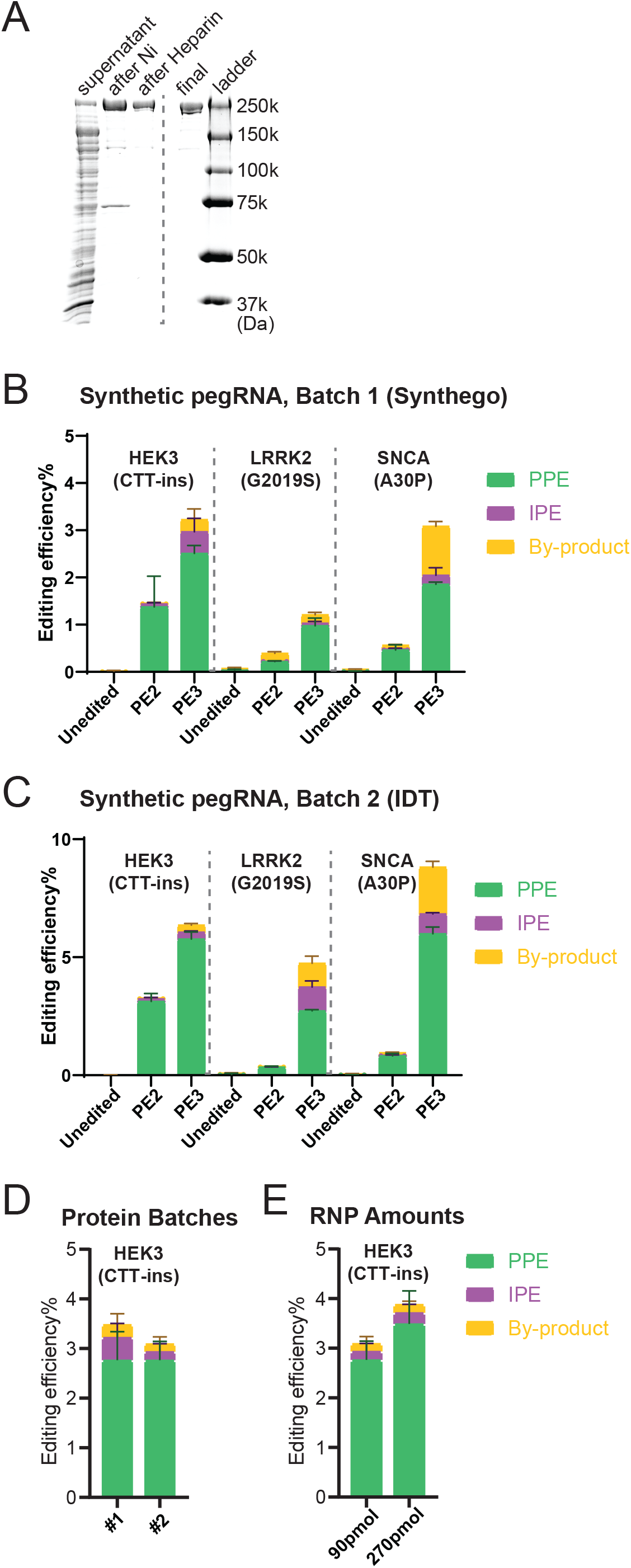
Quality control and parameter testing of RNP-based prime editing. **A.** Coomassie blue staining on at each step during the nCas9-RT protein purification (PE2 version of the prime editor protein as described in Anzalone et al. [2019]). **B.** Bulk prime editing outcomes of RNP-based delivery on three mutations in hESCs with indicated batch 1 of synthetic pegRNA/ngRNAs. A subset of the data shown in figure 2E is analyzed in this panel. N=3. **C.** Bulk prime editing outcomes of RNP-based delivery on three mutations in hESCs with indicated batch 2 of synthetic pegRNA/ngRNAs. A subset of the data shown in figure 2E is analyzed in this panel **D.** Comparison of bulk prime editing outcomes between two batches of purified proteins on *HEK3* (CTT insertion) edits in hESCs. N=3. **E.** Comparison of bulk prime editing outcomes between two doses of RNP on *HEK*3 (CTT insertion) edits in hESCs. N=3. (Error bars indicate the standard deviation [SD], N = number of biological replicates).

**Figure S5.**
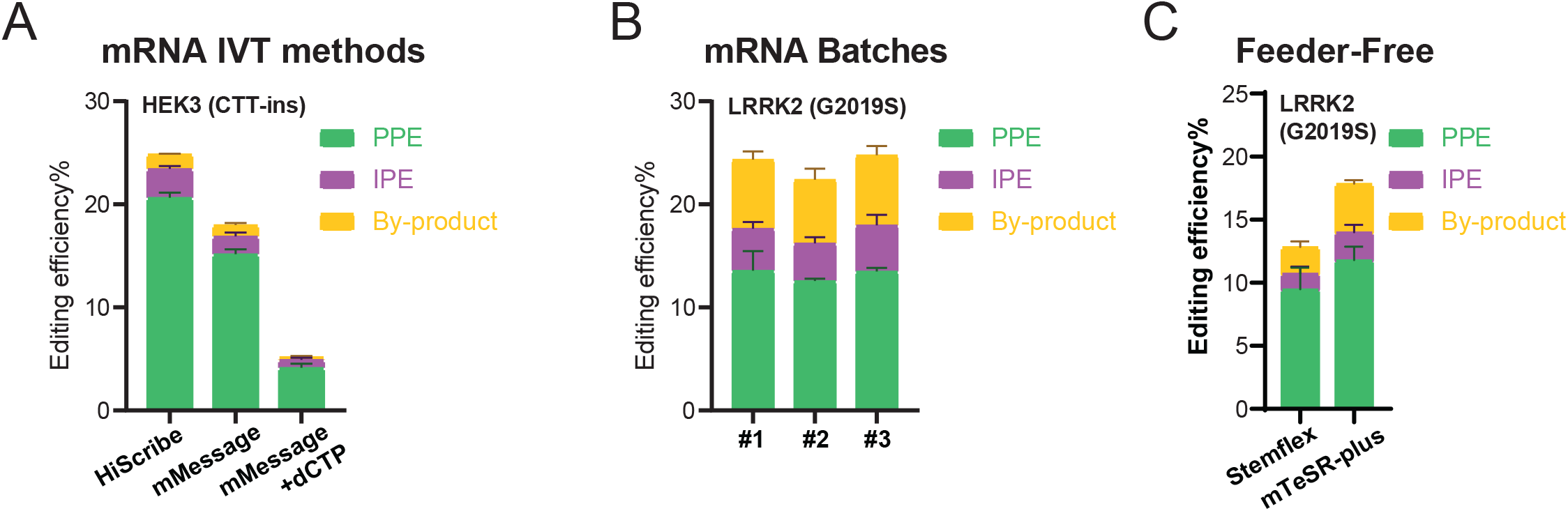
Quality control, parameter testing and prime editing of feeder-free hESC cultures with mRNA-based delivery. **A.** Comparison of bulk prime editing outcomes between three mRNA *in vitro* transcription methods on *HEK3* (CTT insertion) edits in hESCs. N=3. **B.** Comparison of bulk prime editing outcomes between three batches of *in vitro* transcribed mRNA on *HEK3* (CTT insertion) edits in hESCs (HiScribe mRNA *in vitro* transcription). N=3. **C.** Bulk prime editing outcomes on *LRRK2* (G2019S) in hESCs cultured in feeder-free conditions. N=3. (Error bars indicate the standard deviation [SD], N = number of biological replicates).

**Supplemental Table 1:**
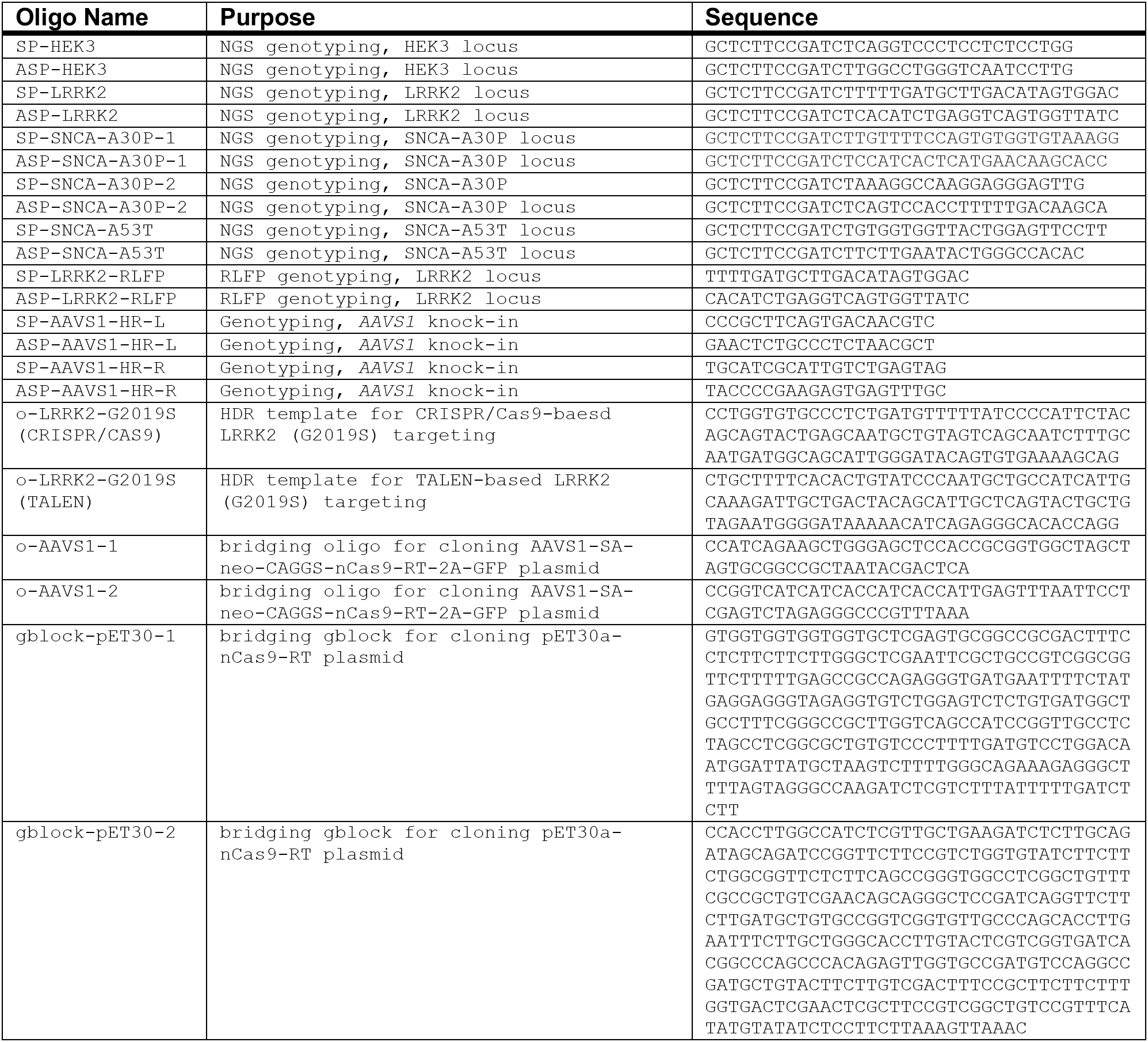

**Supplemental Table 2:**
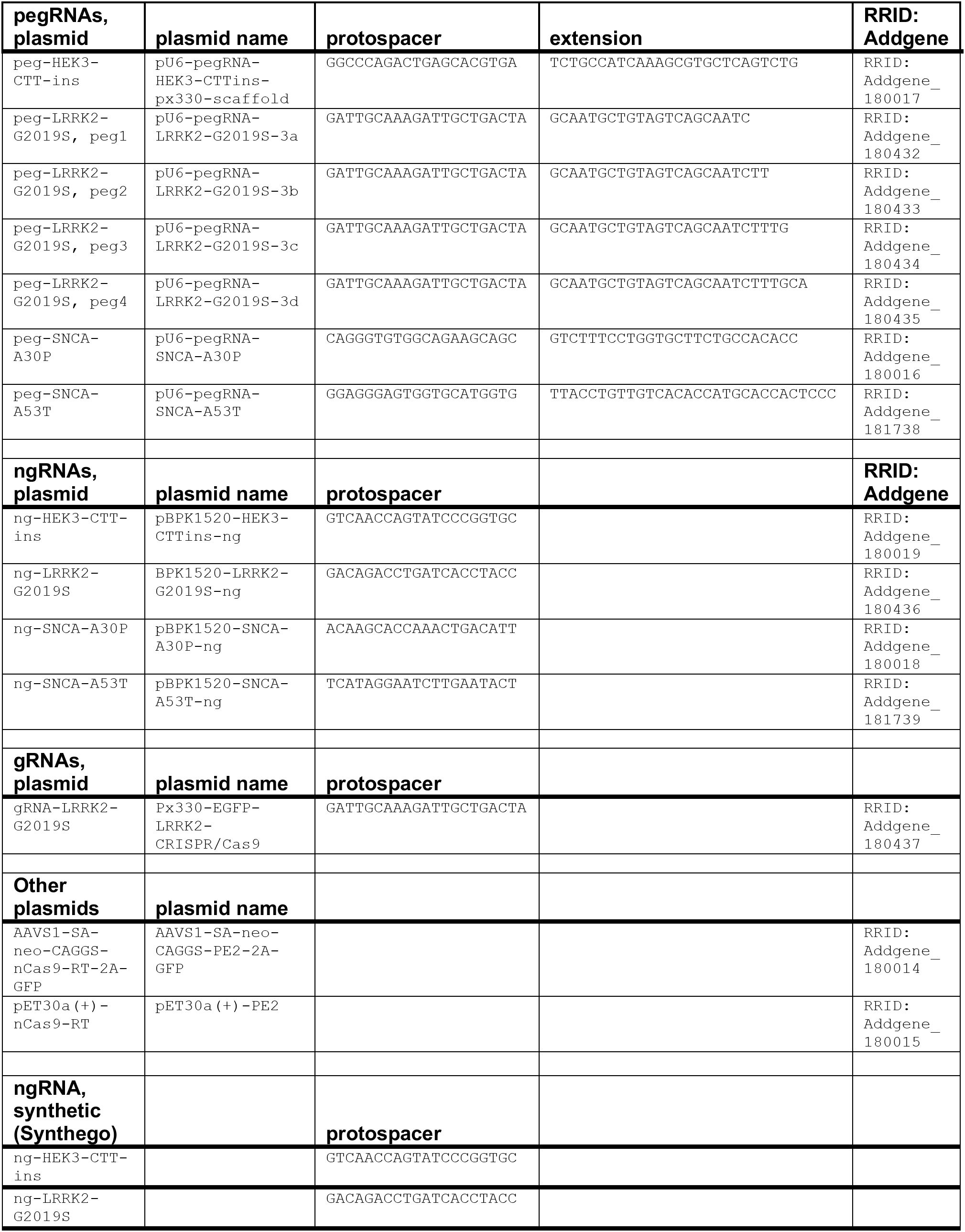
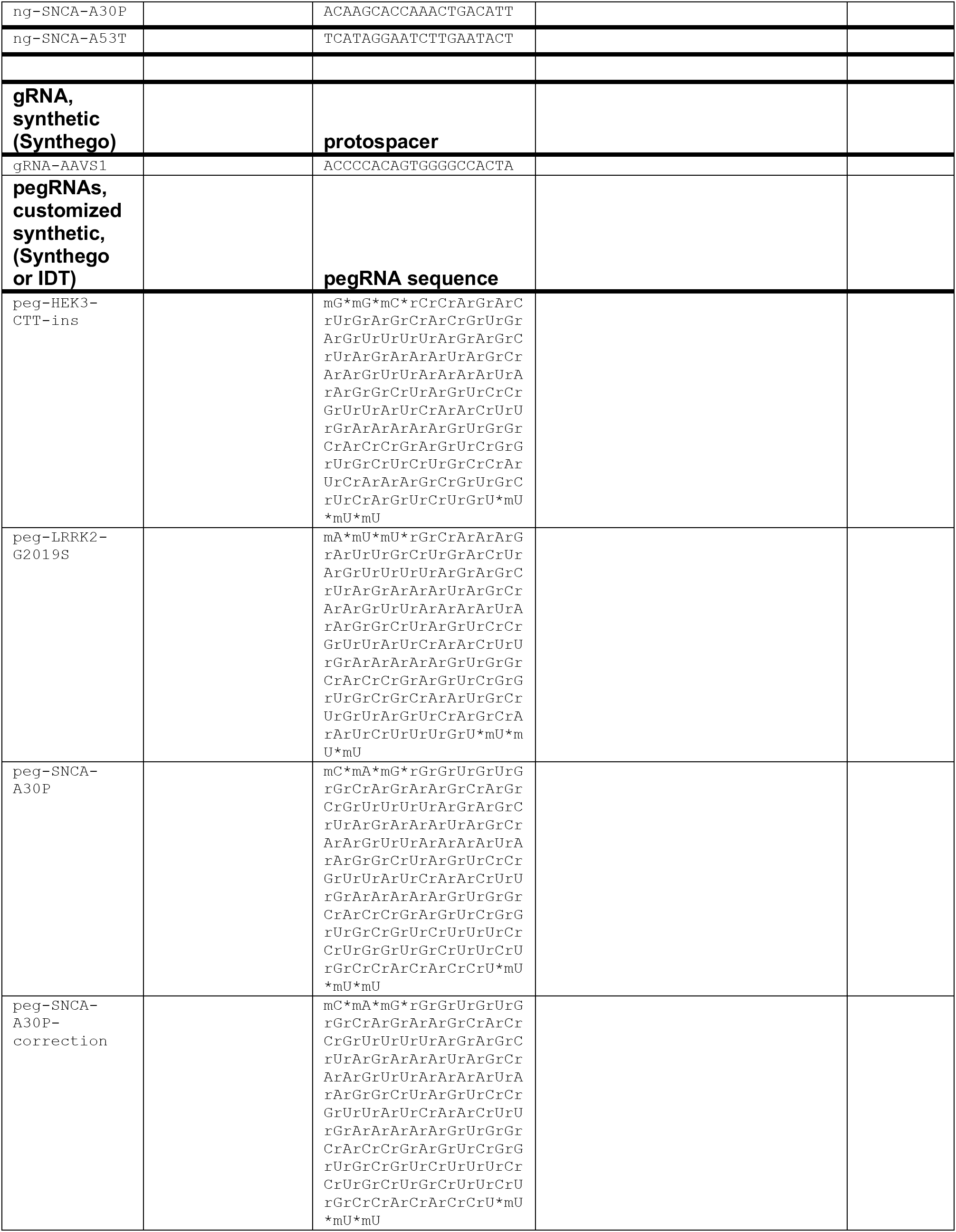

## Notes

### Competing Interest Statement

The authors have declared no competing interest.

